# QuickEd: High-performance exact sequence alignment based on bound-and-align

**DOI:** 10.1101/2024.09.13.612714

**Authors:** Max Doblas, Oscar Lostes-Cazorla, Quim Aguado-Puig, Cristian Iñiguez, Miquel Moreto, Santiago Marco-Sola

## Abstract

**Motivation:** Pairwise sequence alignment is a core component of multiple sequencing-data analysis tools. Recent advancements in sequencing technologies have enabled the generation of longer sequences at a much lower price. Thus, long-read sequencing technologies have become increasingly popular in sequencing-based studies. However, classical sequence analysis algorithms face significant scalability challenges when aligning long sequences. As a result, several heuristic methods have been developed to improve performance at the expense of accuracy, as they often fail to produce the optimal alignment.

**Results:** This paper introduces QuickEd, a sequence alignment algorithm based on a bound-and-align strategy. First, QuickEd effectively bounds the maximum alignment-score using efficient heuristic strategies. Then, QuickEd utilizes this bound to reduce the computations required to produce the optimal alignment. Using QuickEd’s bound-and-align strategy, we reduce *O*(*n*^2^) complexity of traditional dynamic programming algorithms to *O*(*nŝ*), where *n* is the sequence length and *ŝ* is an estimated upper bound of the alignment-score between the sequences. As a result, QuickEd is consistently faster than other state-of-the-art implementations, such as Edlib and BiWFA, achieving performance speedups of 1.6*−*7.3× and 2.1 *−* 2.5×, respectively, aligning long and noisy datasets. In addition, QuickEd maintains a stable memory footprint below 50 MB while aligning sequences up to 1 Mbp.

**Availability:** QuickEd code and documentation are publicly available at https://github.com/maxdoblas/QuickEd.

## 1 Introduction

Pairwise sequence alignment seeks to assess the similarity between two sequences. In bioinformatics, it is a fundamental building block for most analysis tools and pipelines, such as genome assembly (Koren *et al*., 2017; Simpson *et al*., 2009), read mapping (Li, 2018; Marco-Sola et al., 2012; Marco-Sola and Ribeca, 2015) and variant calling (DePristo *et al*., 2011), to name a few.

Traditional pairwise alignment algorithms rely on dynamic programming (DP) (Needleman and Wunsch, 1970; Myers, 1999). These algorithms require computing a matrix proportional to *n* (i.e., length of the sequences to be aligned). Thus, DP-based alignment algorithms usually have a *O*(*n*^2^) complexity in time and memory. Historically, DP-based alignment algorithms have proven effective for aligning relative short sequences (i.e., between 100 and 300 base pairs) like those generated by second-generation sequencing machines. However, their quadratic complexity becomes prohibitively expensive when aligning longer sequences like those generated by third-generation sequencing technologies.

Over the last decades, multiple optimizations have been proposed to boost the performance of classic alignment algorithms. Many optimization strategies are focused on exploiting intra-sequence parallelism to accelerate the DP-matrix computation. For instance, some strategies use data-layout transformations to calculate the DP-matrix along its antidiagonals (Suzuki and Kasahara, 2018; Farrar, 2007). Changing the data access pattern, these optimized algorithms can exploit vector units of modern CPUs to compute several DP-elements in parallel. Other strategies leverage bit-parallel optimizations, where DP-cells are bit-packed in general-purpose registers, enabling computation of several DP-elements using bitwise operations (Myers, 1999; Loving *et al*., 2014). Alternatively, other optimization strategies exploit inter-sequence parallelization by aligning multiple sequences in parallel using the SIMD units of modern processors (Li, 2013; Rognes and Seeberg, 2000). Although most sequence alignment algorithms are typically implemented on general-purpose CPUs, there are notable alternatives specifically developed to run on GPUs (Ahmed *et al*., 2019; Aguado-Puig *et al*., 2023, 2022), FPGAs (Haghi *et al*., 2023, 2021), hardware accelerators (Cali *et al*., 2020; Turakhia *et al*., 2018), and even designed to leverage custom ISA extensions (Doblas *et al*., 2023; Schmidt et al., 2023; López-Villellas et al., 2024; Soria-Pardos et al., 2022). However, the scalability of the algorithms they implement inherently limits these implementations.

Advancements in third-generation sequencing technologies have enabled the production of increasingly longer sequences while increasing production yields and reducing overall costs (Kim *et al*., 2019; Wang *et al*., 2021). Although the accuracy of these technologies has significantly improved (Liu-Wei *et al*., 2024), the sequencing error-rate remains relatively high when producing ultra-long sequences. Therefore, computing the alignment of very long and noisy sequences poses a challenge to traditional DP algorithms, often requiring terabytes of main memory and many CPU hours.

To address this challenge, many algorithms rely on heuristic strategies to improve performance at the expense of accuracy (i.e., approximated algorithms). Commonly used heuristics include banded (Ukkonen, 1985b) and window-based (Turakhia *et al*., 2018) strategies, which accelerate the alignment execution by restricting the solution space explored (i.e., elements computed from the DP-matrix). Other heuristics, such as the X-drop (Suzuki and Kasahara, 2017) and Z-drop (Li, 2018), dynamically prune the solution space and allow early termination by detecting marked drops in the alignment-score. However, restricting the solution space can lead to a loss in accuracy and produce suboptimal alignments, especially when aligning noisy sequences, jeopardizing the accuracy results of the downstream analysis. Hence, we require exact algorithmic strategies that accelerate the alignment computation while generating optimal results (Marco-Sola *et al*., 2023; Groot and Ivanov, 2024; Faust and Hall, 2012).

In contrast, efficient exact alignment algorithms seek to minimize the number of computed DP-elements while capturing the optimal alignment. A classic example of this strategy is the band-doubling technique employed in Edlib (Šošić and Šikić, 2017), which repeats the alignment starting with a narrow band and doubles its size until the optimal alignment is guaranteed to be found within the band. Another example is the wavefront alignment (WFA) algorithm (Marco-Sola *et al*., 2021), which computes the DP-elements in increasing order of score until the optimal alignment is found. Other approaches (Groot and Ivanov, 2024) employ the A^*^ algorithm to guide the computation of the DP-matrix using a battery of heuristics to reduce the solution space explored. While these algorithmic strategies effectively decrease the number of computed DP-elements, they suffer from repeated computations (e.g., repeated band-doubling), additional computational overheads (e.g., computing heuristics), and irregular computation patterns. Compared to classical DP-based algorithms, these approaches require many more computations per each DP-element calculated.

Most of these algorithms perform an uninformed solution space exploration (assuming at first no limit on the alignment score) to avoid missing the optimal alignment. We argue that prior estimation of the alignment-score can significantly reduce the solution space exploration to find the optimal alignment.

In this work, we propose a bound-and-align strategy that first upper-bounds the score of the optimum alignment and subsequently, based on this bound, computes a reduced portion of the DP-matrix using highly efficient alignment strategies. Leveraging the bound-and-align strategy, we present QuickEd, a high-performance sequence alignment algorithm and library. QuickEd consists of two primary components: QuickEd-Bound, which rapidly computes an approximation of the alignment-score using fast and memory-efficient heuristic algorithms, and QuickEd-Align, which exploits high-performance DP-based alignment techniques (with reduced computational cost per cell) to compute a reduced solution space and produce the exact solution (i.e., optimal alignment). Hence, QuickEd can be used for (1) ultra-fast approximation of the sequence alignment (i.e., QuickEd-Bound) or (2) fast and exact sequence alignment (i.e., QuickEd-Bound + QuickEd-Align). In the case of sequence alignment approximation (QuickEd-Bound), we prove that our approach is *O*(*n*) linear in time and *O*(1) constant in memory. In the case of exact sequence alignment (QuickEd-Bound + QuickEd-Align), we prove that our approach reduces the alignment complexity from classic DP-based *O*(*n*^2^) to *O*(*nŝ*), where *n* is the sequence length and *ŝ* is the upper bound of the optimal alignment score.

The rest of the article is structured as follows. Section 2 provides background information to facilitate comprehension of our proposal. Section 3 presents the bound-and-align strategy and introduces QuickEd, a high-performance sequence alignment algorithm based on the bound-and-align strategy. Section 4 shows our proposal’s experimental evaluation and results compared to the state-of-the-art tools and libraries. Finally, Section 5 discusses and summarises the results and contributions of this work.

## 2 Background

### 2.1 Sequence Alignment

Given two sequences and a scoring function (or distance function), the optimal alignment is the sequence of operations (i.e., match, mismatch, insertion, and deletion) that transforms one sequence into the other, maximizing the score (or minimizing the distance).

Given the input sequences, pattern *P* = *p*_0_*p*_1_ … *p*_*n−*1_ and text *T* = *t*_0_*t*_1_ … *t*_*m−*1_, under the edit distance (a.k.a. Levenshtein distance), the optimum edit distance (*Hn,m*) is computed using the DP-recurrence relations shown in Eq 1, where *H*_0,0_ = 0 and *eq*_*i,j*_ = 1 if *p*_*i*_ == *t*_*j*_ and 0 otherwise.

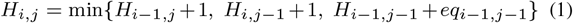

As a result of computing the (*n* + 1) × (*m* + 1) DP-matrix, *Hn,m* contains the optimal distance (or score). The optimal alignment can be backtraced from *H*_*n,m*_ to *H*_0,0_ following the sequence of minimum distance cells that generated the optimum distance.

### 2.2 Bit-parallel Techniques

Bit-parallel techniques emerged in the 80s as an effective strategy to accelerate classic DP-based sequence alignment. These techniques exploit bitwise operations to compute multiple elements of the DP-matrix in parallel. Most importantly, they map exceptionally well to conventional hardware instructions, outperforming other sequence alignment approaches in practice.

The most notable bit-parallel algorithm is the so-called Bit-Parallel Myer’s algorithm (BPM) (Myers, 1999). The BPM algorithm benefits from the observation that differences between adjacent row and column elements in *H* are limited to {−1, 0, +1} values. BPM exploits this property encoding vertical differences (Δ*v*_*i,j*_ = *H*_*i,j*_ *− H*_*i−*1,*j*_) and horizontal differences (Δ*h*_*i,j*_ = *H*_*i,j*_ *−H*_*i,j−*1_) using only (2 ×2) bits per element of the original DP-matrix, irrespectively of the alignment-score. Then, it reformulates the classical DP-equations in terms of differences (Eq. 2).

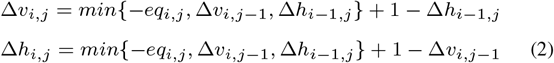

BPM computes the DP-matrix column-wise, packing the DP-elements of each column in a bit-vector. Using bitwise operations, the BPM computes each bit-encoded column using only 17 CPU instructions per character aligned. As a result, the BPM requires to perform 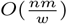computations, where *w* is the machine-word length (usually, *w* = 64). In practice, the BPM outperforms other bit-parallel algorithms (Navarro, 2001), scaling to longer sequences and higher error-rates (Zhang *et al*., 2019). Multiple tools in genome sequence analysis (Šošić and Šikić, 2017; Myers, 2014; Marco-Sola et al., 2012) and hardware accelerators (Doblas *et al*., 2023) have adopted variations of the BPM algorithm.

### 2.3 Windowed Algorithms

Notwithstanding, classic sequence alignment algorithms are intrinsically quadratic. As a result, heuristic methods, like Banded and X-drop, have become increasingly popular. While they are not guaranteed to produce optimal alignments, these methods significantly decrease the time and space requirements by computing only a reduced portion of the DP-matrix. However, even classical heuristics struggle to scale to long sequence alignment as they need to store a prohibitive large portion of the DP-matrix to recover the alignment traceback.

To address this problem, the windowed heuristic introduced in GACT (Turakhia *et al*., 2018) proposes to compute small local alignments within small windows of the DP-matrix and paste them together to produce a global alignment approximation. More specifically, the windowed heuristic starts computing a small window of size *W* × *W* at the bottom-right corner of the DP-matrix. Then, this technique performs a traceback within the window to extract a partial alignment. Afterward, it shifts the window towards the top-left corner of the DP-matrix, leaving an overlap of size *O* with respect to the previous window. This way, the windowed heuristic proceeds by shifting the window, aiming to capture optimal alignment, and pasting all partial alignments together until an end-to-end alignment is retrieved. As a result, this heuristic allows computing a global alignment in linear time, reducing the memory requirements to a single small window of size *O*(*W* ^2^).

The windowed heuristic enables efficient long-sequence alignment computations, thanks to their linear time complexity (with respect to the sequence length) and fixed memory requirements. Moreover, algorithms leveraging this heuristic are well-suited for hardware acceleration because they require a small and predictable amount of memory. However, this heuristic often yields suboptimal alignments, especially aligning noisy and repetitive sequences (Lindegger *et al*., 2023) when gaps are larger than the window size *W*. Thus, it is unsuitable for aligning long and noisy sequences generated by modern sequencers.

Notwithstanding, some state-of-the-art software and hardware-accelerated tools have adopted the windowed heuristic to improve performance at the expense of accuracy. For instance, Scrooge (Lindegger *et al*., 2023) is an edit distance alignment library that implements the windowed heuristic. It computes each window using the Bitap algorithm (Baeza-Yates and Gonnet, 1992), featuring a time complexity of *O*(*W* ^2^*n/*(*W − O*)) and a *O*(*W* ^3^) memory footprint. Similarly, Darwin is a hardware accelerator for gap-affine sequence alignment that utilizes the windowed heuristic to accelerate the classical Smith-Waterman-Gotoh algorithm, achieving a time complexity of *O*(*W* ^2^*n/*(*W − O*)) and a *O*(*W* ^2^) memory footprint.

## 3 QuickEd Algorithm

In this section, we introduce the methods that the QuickEd library implements. First, we present the bound-and-align strategy and QuickEd’s main components (Section 3.1). Then, we present QuickEd’s score-bounding algorithm (QuickEd-Bound, Section 3.2) and QuickEd’s score-bounded alignment algorithm (QuickEd-Align, Section 3.3).

### 3.1 QuickEd’s Bound-and-Align Strategy

Exact alignment algorithms are usually computationally and memory demanding, as they need to explore extensive areas of the DP-matrix.

In contrast, heuristic algorithms are orders of magnitude faster but often fail to produce the optimal alignment. In this work, we aim to combine the performance of heuristic algorithms with the accuracy of exact algorithms using the bound-and-align strategy for efficient exact sequence alignment. The bound-and-align strategy breaks down the alignment process into *bound* and *align* steps (shown in Fig. 1). First, during the *bound* step, an efficient heuristic algorithm computes an upper bound *ŝ* on the optimal alignment-score. In and on itself, a score-bound algorithm can be particularly useful for applications that only require an alignment-score approximation (Xin *et al*., 2015; Alser *et al*., 2019). However, aggressive heuristics algorithms often lose substantial accuracy when aligning noisy sequences. For that, the *align* phase computes the optimal alignment, using the *ŝ* bound to reduce the number of DP-elements computed. The *ŝ* upper bound is continuously used to minimise the region of the DP-matrix explored (i.e., scores that exceed *ŝ*). As a result, the bound-and-align strategy reduces the computational cost of the alignment step from *O*(*n*^2^) to *O*(*nŝ*).

**Fig. 1:**
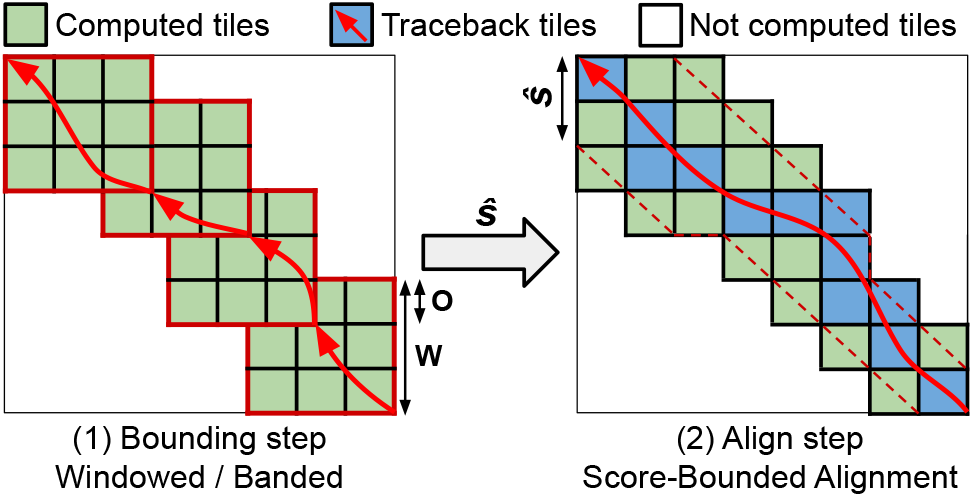
Simplified overview of the bound-and-align strategy implemented in QuickEd. Each square represents a DP-tile (64 × 64 DP-elements).A windowed algorithm computes *ŝ*. (2) A score-bounded alignment algorithm uses *ŝ* continually to reduce the number of DP-tiles computed.

QuickEd is a high-performance sequence alignment algorithm based on the bound-and-align strategy. QuickEd comprises two independent building blocks: QuickEd-Bound and QuickEd-Align (shown in Fig. 2). QuickEd-Bound takes two sequences and produces an accurate score estimation *ŝ* using a cascade of different bounding algorithms. Afterwards, QuickEd-Align aligns the two sequences, exploiting *ŝ* -bound information to reduce the overall DP-cells computed and using a Hirschberg-based implementation to reduce the memory footprint to *O*(*n*); i.e., linear complexity.

**Fig. 2:**
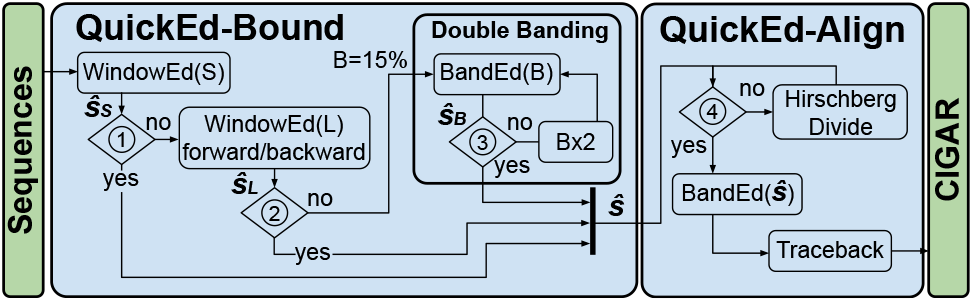
Flow chart of QuickEd’s bound-and-align strategy. Blocks➀-➁ check whether WindowEd(S) and WindowEd(L) had a percentage of HEW lower than a 15% threshold. Block➂checks if *ŝ* _*B*_ *< B*. Block➃ checks if the memory footprint of the alignment (*max*(*n, m*) *· ŝ*) is lower than a configurable threshold.

At the core, QuickEd implements a tiling strategy that divides the entire DP-matrix into square tiles of *T* × *T* elements. This approach offers two key advantages: (1) it allows optimising the tile computation in isolation and tailored to a specific computer architecture, and (2) it offers coarse-grained control over cell computation, which reduces control overhead in the alignment process. In particular, QuickEd’s core relies on the BPM algorithm to efficiently compute 64 × 64 elements tiles. Leveraging this method, every column of 64 DP-elements in a tile is computed using 17 arithmetic-logical instructions. This leads to a 3.76 cell-to-instruction ratio and requires only 2-bit memory per DP-element computed, as only vertical differences need to be stored.

### 3.2 QuickEd-Bound: Score-Bounding Algorithm

Following the bound-and-align strategy, a score-bounding algorithm seeks to produce an upper bound *ŝ* on the optimal alignment-score (*bound* step). To be effective, a bounding algorithm has to (1) run fast and require minimal memory, (2) produce a tight upper bound (or otherwise be able to detect significant deviations from the optimal alignment-score), and (3) exhibit a predictable execution time. Not required to compute the exact optimal alignment-score, a score-bounding algorithm can employ aggressive heuristics, such as the windowed and banded strategies, to be fast and use minimal memory. Moreover, this step is often not required to compute the alignment’s traceback, allowing for a major reduction in memory.

QuickEd-Bound implements a multi-stage cascade of score-bounding algorithms (called *components*), combining instances of WindowEd (Section 3.2.1) and BandEd (Section 3.2.2) for different error tolerances, offering different compromises between accuracy and performance.

#### 3.2.1 WindowEd score-bounding algorithm

WindowEd is a high-performance implementation of the windowed algorithm based on the BPM kernel. The efficiency of the BPM kernel enables efficient scaling to large windows, making it suitable for handling noisy sequences without significantly compromising accuracy.

The window sizes (*W*) and overlap sizes (*O*) used in WindowEd can be configured to different values multiples of the tile size. Depending on the *W* and *O* parameters, window algorithms can suffer from a significant loss in accuracy. For instance, if the optimal sequence alignment contains an indel larger than the window (*W*), the following computed window may completely miss the optimal alignment and substantially overestimate the alignment-score. To mitigate this problem, we introduced a mechanism to detect High-Error Windows (HEWs) as those with an alignment-score higher than a given threshold. We have observed that the number of HEWs in an alignment correlates with an increased probability of missing the optimal alignment. HEWs information is used in the QuickEd-Bound algorithm to determine whether a more accurate score estimation is necessary (e.g., using larger windows).

Based on the BPM and for a CPU word of size *w*, each WindowEd execution takes 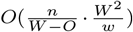 time (i.e., linear on the sequence length).*W −O w* _2_ Regarding memory, each window utilises 2*W* bits of memory for storing the score vertical differences. Since only one window is needed in memory at a time, the memory consumption remains constant and remarkably small for any alignment computation.

#### 3.2.2 BandEd score-bounding algorithm

BandEd is a high-performance implementation of the band-doubling algorithm (Ukkonen, 1985a). This algorithm limits the alignment computation to a band of size *b* around the main diagonal of the DP-matrix under the assumption that the optimal alignment does not stray far from the main diagonal. If the computed score *ŝ* exceeds *b*, the BandEd algorithm may report a sub-optimal alignment-score. In that case, the algorithm repeats the alignment process, doubling the band size (2*b*). The process is repeated until *ŝ ≤b*.

As with previous methods, the BandEd implementation uses a tiling strategy based on the efficient BPM kernel. Additionally, it incorporates a dynamic cut-off optimisation to stop computing tiles once a tile score exceeds *b*. Overall, the BandEd algorithm takes *O*(*nb/w*) time and a constant memory consumption of 2*b* bits, as it only needs to store one DP-column of size *b* using BPM’s differential encoding.

#### 3.2.3 Multi-stage cascade of score-bounding algorithms

Based on WindowEd and BandEd, QuickEd-Bound ultimately implements a multi-stage cascade of score-bounding components, initially using fast components (for highly similar sequences) and then, if necessary, switching to slower but more accurate ones (for highly divergent sequences). Figure 2 (left side) shows QuickEd-Bound’s workflow. Its design comprises three components connected in cascade: WindowEd(S), WindowEd(L), and BandEd(band size). Note that all components compute only the alignment-score, reducing the required memory footprint to a constant value.

The first component of the bounding algorithm, WindowEd(S), is a vectorized implementation of WindowEd that computes the small windows of size *W* = 128 and *O* = 64. In practice, WindowEd(S) is fast and accurate in estimating the alignment-score of highly similar sequences (e.g., Illumina and PacBio HiFi sequences).

The second component, WindowEd(L), uses a bigger WindowEd of size *W* = 640 and *O* = 128. In this case, WindowEd(L) is executed twice, once aligning the sequences forward and then backwards. This strategy seeks to address cases where a large gap in the alignment makes the windowed algorithm miss the optimal path and significantly overestimate the alignment-score. In practice, this component improves the alignment-score estimation on dissimilar sequences while still being fast and efficient due to its linear dependency on the sequence length.

The third and last component, BandEd(B), uses the band-doubling algorithm (Ukkonen, 1985b) to compute the optimal alignment-score when all the other components have failed to produce an accurate estimation. QuickEd’s band-doubling technique starts with a band size *b* = 0.15*n* (i.e., assuming a maximum error-rate of 15%). It doubles the band size *b* until the alignment-score bound *ŝ* is lower than the selected band ➂, ensuring that the optimal alignment-score is found. In practice, this component is the slowest but produces the exact alignment-score, even in the presence of large gaps and very noisy regions.

After each component’s execution, QuickEd-Bound evaluates the number of HEWs found during the windowed alignment ➀-➁) to determine whether a component produces a tight alignment-score bound. If more than 15% of the computed alignment windows exceed 40% error-rate (i.e., less than 60% local similarity), the algorithm proceeds to bound using the next component in the cascade. Otherwise, the algorithm considers the alignment-score bound tight or close to the exact alignment-score and moves on to the alignment step (QuickEd-Align). Unlike QuickEd-Bound, which seeks to compute a close approximation of the alignment-score, QuickEd-Align always guarantees to find the exact alignment solution. A detailed analysis of the parameter space exploration (i.e., window-size and overlap sizes selection) for each component can be found in Supplementary S3.

### 3.3 QuickEd-Align: Score-Bounded Alignment Algorithm

Following the bound-and-align strategy, we seek an exact alignment algorithm that is guaranteed to find the optimum alignment (*align* step). To be effective, it should (1) exploit the *ŝ* bound to reduce the number of DP-elements computed, (2) require minimum control flow to select which DP-elements to compute, and (3) minimise memory requirements usage to scale up with long sequence alignment.

Note that various strategies can exploit the *ŝ* bound to reduce the number of DP-elements computed. However, a dynamic cut-off (Myers, 1999) is often very effective and requires minimum additional computations. In contrast, other cut-off strategies, like A^*^-based algorithms (Groot and Ivanov, 2024; Hadlock, 1988), can introduce substantial overheads, limit parallelism, and require extra memory (e.g., state priority queues and indexing data structures). Yet, combining a dynamic cut-off with Quicked’s underlying tiling strategy enables per-tile cutoff evaluation, reducing the cutoff’s overheads even more.

Therefore, QuickEd-Align implements a per-tile dynamic cutoff using a prior *ŝ* estimation to reduce the number of DP-elements required to compute the optimal alignment between two sequences. This strategy aims to selectively compute DP-elements with a score *s ≤ ŝ*. Specifically, an *H*_*i,j*_ DP-element may belong to an alignment path of score *ŝ* (or lower) if *H*_*i,j*_ + *d*_*k*_(*i, j*) *≤ ŝ*, where *d*_*k*_(*i, j*) = |(*j − i*) *−* (*m − n*)| is the minimum distance from a DP-element to the target diagonal (i.e., diagonal where the alignment ends). Since QuickEd-Align uses a tiling strategy, the cut-off optimisation operates at the DP-tile level. Hence, an entire tile must be computed if, at least, one DP-element within the tile may belong to an alignment path of score *ŝ*. Then, for each tile, QuickEd only needs to examine the DP-element closest to the main diagonal, which can be (a) the bottom-left DP-element for tiles above the target diagonal or (b) the upper-right DP-element for tiles below the target diagonal. Since the cut-off uses an upper bound of the optimal score, QuickEd-Align is guaranteed to find the optimal alignment. Ultimately, QuickEd-Align computes *O*(*nŝ*) DP-elements, resulting in a time complexity of *O*(*nŝ/w*).

#### 3.3.1 Divide-and-conquer algorithm

A direct implementation of QuickEd-Align requires storing all the computed DP-elements to allow tracing-back the optimal alignment afterwards. Using BPM’s differential encoding, this implementation would use a *O*(2*nŝ*) bits of memory. Despite achieving a lower memory bound than *O*(*nm*) DP-based algorithms, aligning long and noisy sequences would lead to substantial memory consumption. When the *O*(*nŝ*) bound anticipates the need for a large memory footprint, QuickEd switches to a Hirscherg-based implementation of QuickEd-Align.

Hirschberg’s technique is used to achieve linear space complexity, and it has been implemented widely across various alignment libraries and tools (Marco-Sola *et al*., 2023; Myers, 1986; Šošić and Šikić, 2017). Hirschberg’s key idea is to use a divide-and-conquer strategy that computes the alignment from both ends of the sequences (i.e., one alignment forward and another backwards), progressing until they converge at the midpoint of the DP-matrix. This way, Hirschberg’s technique can determine a breakpoint where the forward and backwards alignments meet optimally (i.e., DP-element contained in the optimal alignment path), identifying a split point for further problem subdivision. This process is repeated until the full alignment is computed.

It is important to note that each alignment must only store one column at a time, ensuring that only linear space is used during the process.

Moreover, each breakpoint determines the optimal alignment-score *s*\* of each subproblem, allowing QuickEd to use it as the perfect cutoff bound (*ŝ* = *s*\*) and fall back to the regular QuickEd-Align whenever the expected memory usage drops below a configurable threshold (e.g., 16MB matching the last level cache sizes of modern processors).

Fig. 3 shows a graphical example of QuickEd-Align computing a pairwise alignment using Hirschberg’s technique and constrained by a specific bounding score. During the first iteration (Figure 3➀), Hirschberg’s technique is used since the expected memory footprint is higher than the threshold. Therefore, the alignment is computed forward and backwards until reaching the midpoint of the DP-matrix. This way, the algorithm computes the optimal breakpoint (with score *s*\*), divides the alignment into two smaller sub-problems, and adjusts the bounding score using *s*\* to reduce computations in the following iterations. The algorithm proceeds to compute the optimal breakpoints of the two sub-problems (Fig. 3➁) and divide the problem into smaller subproblems, reducing the alignment-score bound used on each iteration. The algorithm iterates until the alignment problems require a small memory footprint and can be computed using the regular QuickEd-Align (Fig. 3.©).

**Fig. 3:**
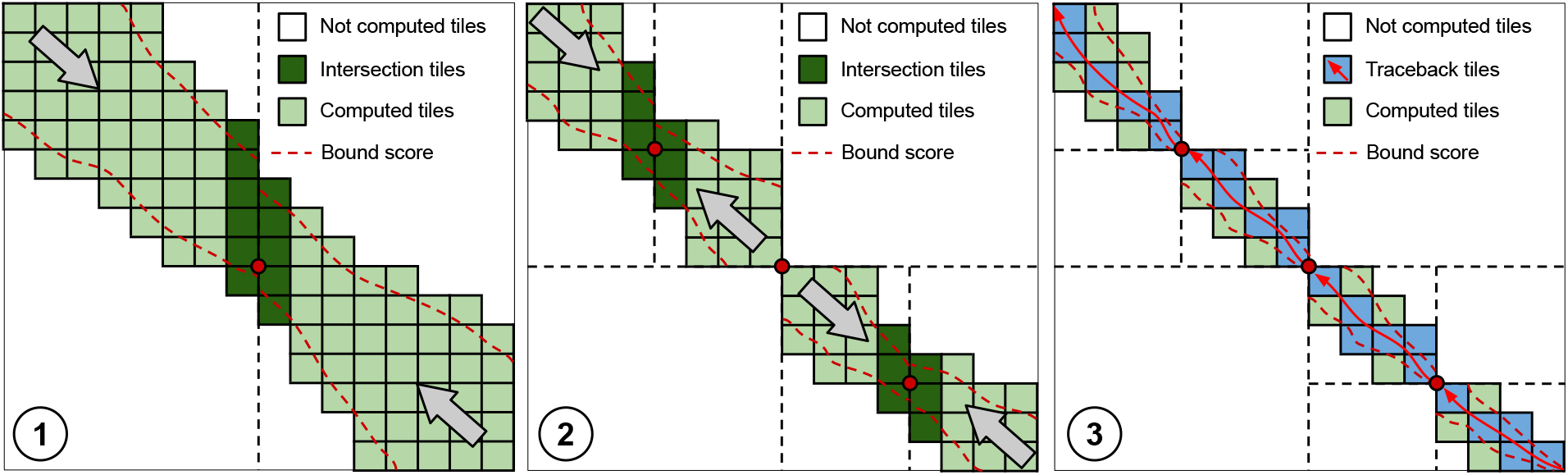
Overview of QuickEd-Align’s algorithm using a *ŝ* score-bound cut-off and Hirschberg technique. Each square represents a computed 64×64 tile (green). Iterations➀ and ➁ divide the alignment into smaller sub-alignments by computing an optimal breakpoint (red dot), storing only a DP-column (dark green). For iteration➂, sub-alignments are sufficiently small to use regular QuickEd-Align and generate the full alignment (pale blue).

## 4 Results

We implemented the QuickEd library in C. The code and the scripts required to reproduce the experimental results presented in this section are publicly available at https://github.com/maxdoblas/QuickEd.

### 4.1 Experimental Setup

We evaluated the performance of QuickEd compared to other state-of-the-art sequence alignment libraries. First, in Section 4.2, we evaluate QuickEd for alignment-score approximation (i.e., alignment-score bounding) and compare it against other approximated and heuristic algorithms: Scrooge (Lindegger *et al*., 2023), the banded heuristic, and windowed heuristic for different parameter configurations. Second, in Section 4.3, we evaluate QuickEd for exact sequence alignment and compare it against other state-of-the-art aligners implementations: Edlib (Šošić and Šikić, 2017), KSW2 (Li, 2018), A^*^PA (Groot and Ivanov, 2024), WFA (Marco-Sola *et al*., 2021), and BiWFA (Marco-Sola *et al*., 2023). We excluded from the evaluation other libraries such as Parasail (Daily, 2016) and SeqAn (Döring *et al*., 2008) as they were not designed to align very long and noisy sequences and failed to complete the executions.

For the evaluation, we use simulated and real datasets. For the evaluation using simulated datasets (see Supplementary S4), we generated datasets of different sequence lengths (i.e., 10K, 100K, 500K, and 1M bases) for a total of 100 million bp, randomly distributed error-rates (i.e., *e* =1%, 5%, 10%, and 20%) using the same methodology used in (Marco-Sola *et al*., 2021; Groot and Ivanov, 2024). For the real datasets, we selected six representative datasets from Illumina (Illumina 250), PacBio (PacBio HiFi), and Oxford Nanopore (ONT UltraLong, ONT PromethION, ONT MiniION-a, and ONT MiniION-b). All datasets are publicly available (see Supplementary S1). For Scrooge executions, we had to modify the input datasets, replacing all ‘N’ characters with ‘A’, as Scrooge only allows {*A, C, G, T*} characters. All executions were performed on an Intel Xeon Platinum 8160 CPU @2.10GHz node equipped with 96GB of DRAM.

### 4.2 Evaluation of Approximated Algorithms

In this section, we evaluate different approximated algorithms for alignment-score bounding in terms of time performance and accuracy. In particular, we evaluate our proposal (QuickEd-Bound) against Scrooge (state-of-the-art approximate sequence alignment algorithm), two instances of WindowEd (i.e., WindowEd(S) and WindowEd(L)), and BandEd using a band size of 15% the sequence length.

To evaluate the accuracy of the results, we calculate the Normalized Score Deviation (NSD). The NSD represents the average alignment-score deviation produced by an approximated algorithm compared to the optimal score, which an exact algorithm would obtain. The NSD is computed as ((*ŝ − s*_*opt*_)*/s*_*opt*_)*/N* where *ŝ* is the alignment-score computed by the approximated algorithm, *s*_*opt*_ is the optimal alignment-score, and *N* is the number of paired sequences in the dataset. Within QuickEd’s bound-and-align framework, the NSD also depicts the additional computations that the align step will perform due to overestimating the alignment-score during the bound step.

Table 1 shows time and NSD for all the approximated algorithms evaluated. Overall, we observe that Scrooge is limited by its smaller window size (W=64) and consistently obtains a worse NSD than other methods. In particular, WindowEd(S) achieves a 2.3 *−* 21.3× NSD improvement while executing 1.6 *−* 11.7× faster than Scrooge. In the case of WindowEd, using the BPM kernel enables faster computation of larger windows (W=128) while reducing memory usage and improving the accuracy of the results (i.e., smaller NSD).

**Table 1.** Time (in seconds) and Normalized Score Deviation (NSD, in %) for various approximated algorithms aligning real datasets. ^†^ For Scrooge executions, we modify the input datasets, replacing all ‘N’ characters with ‘A’, as Scrooge only allows {*A, C, G, T*} characters.

|  | Scrooge<sup>†</sup> |  | WindowEd(S) |  | WindowEd(L) |  | BandEd(15%) |  | QuickEd-Bound |  |
| --- | --- | --- | --- | --- | --- | --- | --- | --- | --- | --- |
|  | Time (s) | NSD (%) | Time (s) | NSD (%) | Time (s) | NSD (%) | Time (s) | NSD (%) | Time (s) | NSD (%) |
| Illumina 250bp | 56.0 | 0.3 | 34.8 | 0.0 | 66.6 | 0.0 | 69.0 | 0.0 | 43.2 | 0.0 |
| PacBio HiFi | 370.8 | 505.3 | 196.8 | 30.4 | 882.0 | 5.7 | 2400.6 | 0.0 | 312.6 | 5.2 |
| ONT UltraLong | 372.0 | 209.8 | 45.4 | 36.7 | 154.2 | 8.8 | 3302.4 | 10.9 | 549.6 | 1.0 |
| ONT PromethION | 39.4 | 751.8 | 3.9 | 35.3 | 16.2 | 2.5 | 718.2 | 1.4 | 6.0 | 1.5 |
| ONT MiniION-a | 6.3 | 1209.2 | 0.5 | 94.5 | 2.1 | 16.8 | 316.2 | 1.4 | 7.2 | 6.1 |
| ONT MiniION-b | 6.7 | 936.5 | 0.7 | 402.8 | 2.3 | 144.8 | 348.0 | 1.4 | 58.8 | 15.5 |

Despite this improvement, WindowEd(S)’s NSD results can be significantly improved, especially when aligning ONT datasets. Using larger windows (W=640), WindowEd(L) obtains an average 5.5× NSD improvement while being 3.5× slower than WindowEd(S). Nonetheless, WindowEd(L) still remains 2.0× faster on average than Scrooge, even though it computes 10× larger windows.

Moreover, results on Table 1 indicate that BandEd(15%) is the most accurate alignment-score estimator, achieving a lower NSD in most cases except with ONT-UltraLong. However, BandEd(15%) execution time increases up to 153× compared to WindowEd(S) when aligning the ONT datasets, making it computationally very expensive.

Ultimately, these results support the claim that a combination of WindowEd(S), WindowEd(L), and BandEd (i.e., QuickEd-Bound) is the most sensible design towards a compromise between accuracy and performance. In fact, results in Table 1 show that QuickEd-Bound can achieve similar NSD as BandEd(15%) while requiring a similar execution time to the WindowEd approaches. In practice, QuickEd-Bound cascade design can perform on par with the WindowEd methods while producing accurate results comparable to those of BandEd for estimating the alignment-scores even when aligning long and noisy sequences.

More in detail, Fig. 4 displays the cumulative alignment-score count for the different approximated algorithms aligning real datasets. Note that the approximated alignment-score is always greater or equal to the optimal alignment-score (*ŝ < s*\*). Hence, the curve generated by an exact algorithm (e.g., Edlib as baseline) helps to identify alignment-score estimation deviations produced by approximated algorithms. This way, the cumulative alignment-score curves show that WindowEd(S) achieves more accurate score estimations than Scrooge, particularly when aligning ONT datasets. Moreover, these results show the necessity of using BandEd to align noisy sequences, as windowed approaches can produce poor accuracy results. Ultimately, these results demonstrate that QuickEd-Bound can efficiently produce accurate alignment-score approximations, significantly outperforming other approximated methods.

**Fig. 4:**
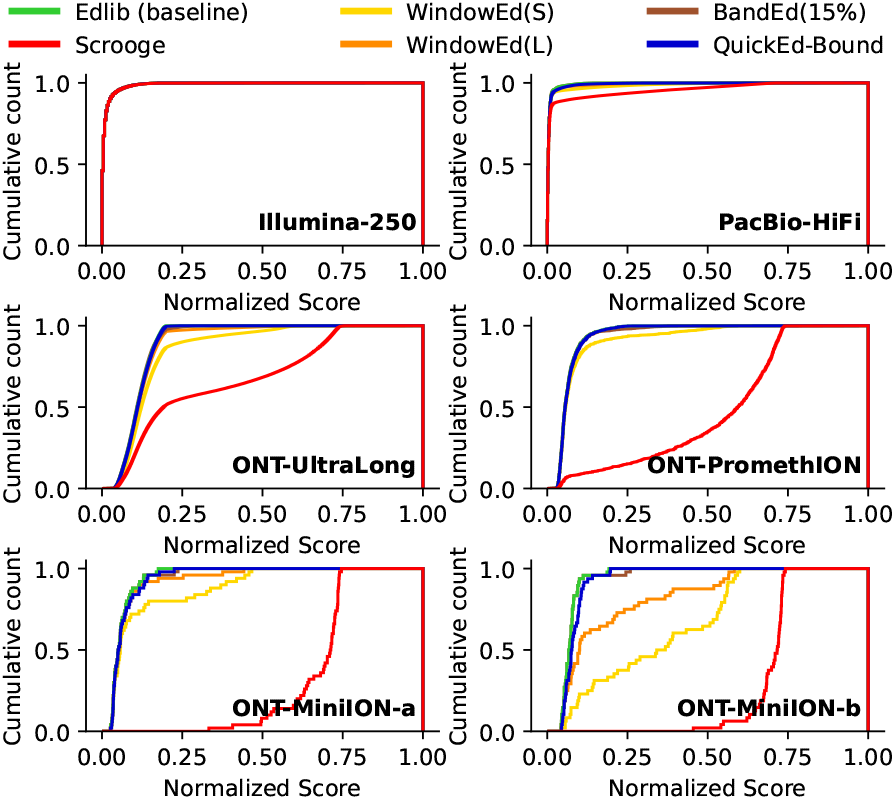
Cumulative alignment-score distribution curves for various approximated algorithms aligning real datasets.

### 4.3 Evaluation of Exact Alignment Algorithms

In this section, we evaluate the performance of exact sequence alignment algorithms. In contrast to approximated algorithms from the previous section, these algorithms are always guaranteed to produce optimal results (i.e., 100% recall). In the following, we evaluate QuickEd against Edlib, KSW2, A^*^PA, WFA, and BiWFA.

Table 2 shows performance results (time and memory usage) aligning real datasets. We observe that QuickEd is 3.1 *−* 7.3× faster than Edlib and 9.1 *−* 22.7× faster than A^*^PA when aligning high-quality sequences (i.e., Illumina and PacBio datasets). In contrast, WFA-based implementations, which exploit sequences’ similarities to accelerate the alignment computation, perform 7.1 *−* 16.3× faster than QuickEd. However, when aligning long and noisy ONT datasets, QuickEd consistently outperforms all other implementations. In particular, QuickEd performs 1.6 *−*2.5× faster than Edlib, 2.1*−*2.5× faster than WFA-based implementations, and up to 78.3× than A^*^PA (considering only successful A^*^PA executions that do not run out of memory).

**Table 2.** Time (in seconds) and memory consumption (in megabytes) for the different exact alignment tools aligning different real datasets. Executions taking more than 48 hours are marketed as “to” (time out) and executions as run out of memory as “oom”.

|  | Time (s) |  |  |  |  |  | Memory (MB) |  |  |  |  |  |
| --- | --- | --- | --- | --- | --- | --- | --- | --- | --- | --- | --- | --- |
|  | KSW2 | WFA | BiWFA | A*PA<sup>†</sup> | Edlib | QuickEd | KSW2 | WFA | BiWFA | A*PA<sup>†</sup> | Edlib | QuickEd |
| Illumina 250bp | 1185.0 | 7.6 | 15.8 | 1124.4 | 403.2 | 124.2 | 1.1 | 1.1 | 1.1 | 1.8 | 18.8 | 18.6 |
| PacBio HiFi | to | 113.4 | 153.6 | 18140.4 | 5893.8 | 800.4 | to | 85.9 | 3.0 | 246.8 | 20.5 | 40.6 |
| ONT UltraLong | oom | oom | 7075.8 | oom | 5998.8 | 2795.4 | oom | oom | 37.0 | oom | 29.0 | 48.2 |
| ONT PromethION | oom | 696.6 | 880.8 | 27288.6 | 884.4 | 346.2 | oom | 10299.9 | 12.6 | 52279.0 | 21.8 | 40.9 |
| ONT MiniION-a | oom | 316.2 | 366.0 | oom | 322.8 | 159.0 | oom | 47169.3 | 24.3 | oom | 25.5 | 43.4 |
| ONT MiniION-b | oom | oom | 547.2 | oom | 431.4 | 265.2 | oom | oom | 20.8 | oom | 26.5 | 47.6 |

Regarding memory, QuickEd shows very low memory consumption, aligning both short and long datasets, comparable to BiWFA and Edlib, which are known for their minimal memory footprint. In fact, these three libraries implement variations of Hirschberg’s algorithm to drastically reduce the memory footprint. In contrast, KSW2, the original WFA, and A^*^PA require excessive amounts of memory, often running out of memory and failing, which strongly limits their applicability to long and noisy sequence alignment.

Additionally, Table S2 shows performance results (i.e., execution time and memory consumption) using simulated datasets. Overall, these results indicate that QuickEd is generally faster than other methods. In particular, QuickEd consistently outperforms Edlib, being 2.2 *−* 9.7× faster. Note that this performance improvement is particularly notable when aligning short sequences. Compared to KSW2, QuickEd is 9.5× faster aligning Illumina short reads.Similarly, QuickEd demonstrates to be faster than A^*^PA aligning short or very low-error sequences because A^*^PA incurs computation overhead due to the random computation of DP-elements. Moreover, QuickEd outperforms A^*^PA in aligning complex sequences (high divergence, sequences with long indels, or sequences with repetitive regions), since A^*^PA exhibits quadratic time complexity in such cases.

Compared to WFA-based implementation, Quicked is 1.41× faster on average. However, aligning short and similar sequences WFA-based implementations perform better. This is most likely due to QuickEd’s smallest band (set to a minimum of 192 DP-elements or, equivalently, 3 DP-tiles), which limits the effectiveness of the cut-off optimization in such scenarios. In particular, QuickEd outperforms both WFA-based implementations aligning long and noisy sequences, achieving speedups up to 2.36×.

## 5 Discussion

Pairwise sequence alignment remains a core performance-critical component of many bioinformatics tools and pipelines. Modern sequencing technologies have enabled the production of long sequences. However, long sequence analysis poses a critical challenge to the scalability of classic pairwise alignment algorithms in time and memory. Recently proposed methods advocate for aggressive heuristics that prove to be fast and memory efficient at the expense of significantly deviating from the optimal solution, which limits their applicability in real-world applications. On the other hand, novel exact alignment methods, like those based on A^*^ search, effectively reduce the number of DP-elements to compute but introduce a high computational cost per DP-element computed. Furthermore, memory usage is often overlooked, and many exact algorithms require a prohibitive amount of memory when aligning long and noisy reads.

This work presents QuickEd, a high-performance library for approximate and exact sequence alignment based on bound-and-align. Following the bound-and-align strategy, QuickEd can make informed decisions for selecting the most suitable alignment algorithm for each input pair of sequences depending on its characteristics (e.g., length, nominal error, and error distribution). For that, QuickEd mixes efficient heuristic (QuickEd-Bound) and exact methods (QuickEd-Align) to reduce alignment computations while producing optimal results. Using the bound-and-align strategy, QuickEd improves *O*(*n*^2^) time complexity of the classical DP algorithms to *O*(*nŝ*), where *ŝ* is an estimated upper bound on the alignment-score. More importantly, we argue that it is highly effective to leverage the efficiency of heuristic methods to boost exact alignment algorithms and produce optimal results. In particular, QuickEd-Bound employs various heuristic methods to help reduce the computations and complexity of QuickEd-Align to produce optimal results, while enabling hardware-friendly optimisations like bit-parallel computations and tiling.

We evaluated QuickEd against other state-of-the-art implementations to demonstrate the performance benefits of the bound-and-align approach. For approximating the alignment-score, we show that QuickEd-Bound outperforms other state-of-the-art approximate aligners, like Scrooge, achieving on average 1.7× higher throughput while obtaining on average 176.6× lower NSD. For exact sequence alignment, we demonstrate that QuickEd consistently outperforms by 1.6 *−* 2.5× other state-of-the-art exact alignment libraries, including Edlib, A^*^, BiWFA, and KSW2, when aligning long and noisy sequences produced by ONT modern sequencers. In particular, QuickEd is consistently faster than Edlib and BiWFA, achieving performance speedups of 1.6 *−* 7.3× and2.1 *−* 2.5×, respectively, aligning long and noisy datasets. In addition,QuickEd maintains a stable memory footprint below 50 MB, even aligning sequences up to 1 Mbp with a 10% error-rate.

Genomics and bioinformatics methods will continue to rely on pairwise alignment as a core component. QuickEd’s library, based on the bound-and-align strategy, offers a practical and efficient method for approximate and exact sequence alignment of long and noisy sequences. We expect QuickEd to pave the way for high-performance sequence analysis tools, enabling efficient and hardware-friendly implementations that scale with future advances in genome analyses and sequencing technologies.

## Acknowledgements

The authors would like to thank Christopher Batten for his invaluable and insightful comments during many fruitful discussions.

