## Supplementary material for "QuickEd: High-performance exact sequence alignment based on bound-and-align"

#### S1 Experimental Datasets

To evaluate the QuickEd library, we used real datasets produced by Illumina, PacBio, and Oxford Nanopore technologies. Table S1 shows each dataset's number of sequences, minimum, average, and maximum sequence length and error rate. Illumina 250 dataset was obtained from NIST's Genome in a Bottle (GIAB) project and can be found at `https://github.com/genome-in-a-bottle/giab_data_indexes`. PacBio HiFi dataset was obtained from PrecisionFDA Truth Challenge V2 and can be found at `https://precision.fda.gov/challenges/10`. ONT UltraLong dataset, aligned against the v1.1 CHM13 assembly, was obtained from Bowden *et al.* (2019). ONT PromthION dataset was obtained from the Human Pangenome Reference Consortium (Miga and Wang (2021)). ONT MiniION-a dataset contains sequences produced by Oxford Nanopore MiniION and was obtained from `https://github.com/pairwise-alignment/pa-bench/releases/download/datasets/ont-500k.zip`. All sequences are longer than 500kbps. MiniION-b dataset contains sequences produced by Oxford Nanopore MiniION from Bowden *et al.* (2019) aligned against the v1.1 CHM13 assembly. All sequences are longer than 500kbps and contain genetic variation and large gaps.

More in detail, Figure S1 shows sequence length and error distribution for each real dataset.

Table S1: Properties of real datasets used in the experimental evaluation.

| Dataset | No. pairs | Seq. Length (bps) | | | Error rate (%) | | |
|---|---|---|---|---|---|---|---|
| | | min | avg | max | min | avg | max |
| Illumina 250 | 100M | 140 | 248 | 275 | 0.0 | 1.0 | 20.0 |
| PacBio HiFi | 10M | 201 | 12,847 | 24,640 | 0.0 | 0.6 | 20.0 |
| ONT UltraLong | 100k | 125 | 21331 | 1,053,101 | 0.3 | 11.3 | 20 |
| ONT PromethION | 1312 | 14,749 | 174,219 | 309,277 | 2.6 | 6.7 | 33.2 |
| ONT MiniION-a | 50 | 502,992 | 597,582 | 848,888 | 2.7 | 6.1 | 17.6 |
| ONT MiniION-b | 48 | 503,480 | 635,931 | 1,053,101 | 4.4 | 7.4 | 19.2 |

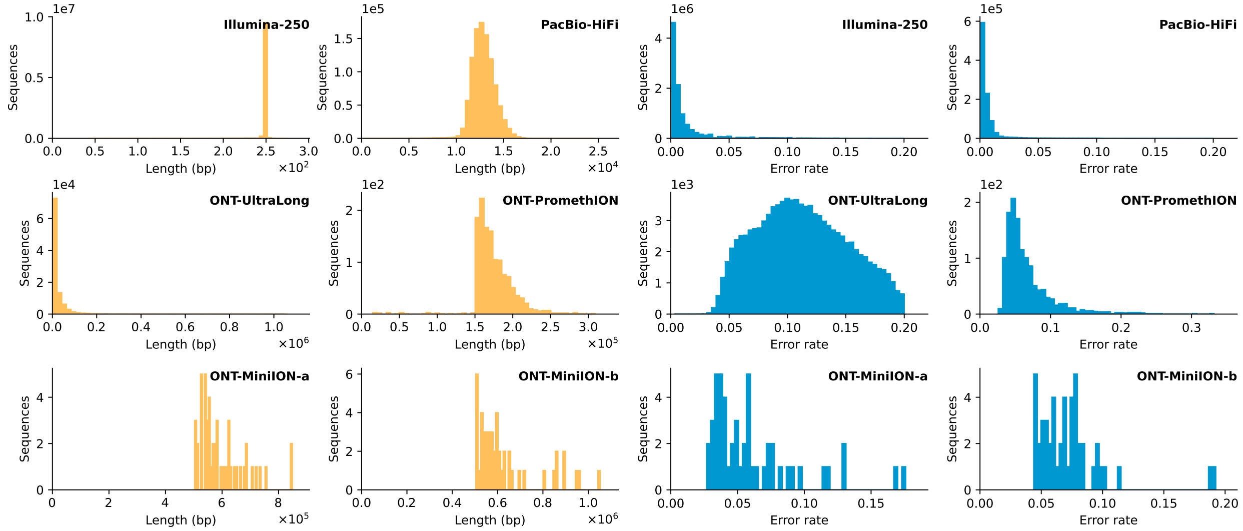


Figure S1: Sequence length (yellow) and error distribution (blue) of all real datasets used in the experimental evaluation.

### S2 QuickEd Time Analysis

QuickEd has two main building blocks: a score-bounding algorithm (bound step) and a score-bounded alignment algorithm (align step). Given a sequence pair $p \in P$, from a set of input sequence pairs $P$, the score-bounding algorithm approximates the alignment-score ($\hat{s}_p$). This estimation may be sub-optimal but serves as a guide to narrow down the alignment search space during the score-bounded align step, reducing the overall execution time.

$$T = \sum_{p \in P} T_B(p) + \sum_{p \in P} T_A(p, \hat{s}_p) \tag{1}$$

The overall execution time ($T$) is the aggregate of the score-bounding time ($T_B$) and the score-bounded alignment time ($T_A$), using $\hat{s}_p$ bound, as shown in Equation 1. Generally, more precise score-bounding algorithms tend to be more time-consuming. However, the execution time of score-bounded alignment algorithms depends on the alignment-score estimation provided by the bounding step. In particular, a score-bounded alignment algorithm becomes more costly if the estimation provided by the bound step is less accurate. Thus, an important trade-off exists between the precision attained during the bound step and the overall execution time (i.e., the sum of the bound and align execution time).

#### S2.1 Quicked's Cascade of Score-Bounding Components

Designing a single score-bounding algorithm that simultaneously delivers good speed and accuracy across all types of sequences is challenging. A more practical approach involves implementing a cascade of multiple score-bounding algorithms (called components), each tailored to handle sequences of different characteristics. This score-bounding is structured hierarchically, progressing from faster, less accurate components to slower, more precise ones. Figure S2 shows a $n$-step score-bounding algorithm with $n$ different components $(B_1, B_2, ..., B_n)$ connected in cascade. The initial component ($B_1$) should be a fast and simple score-bounding algorithm able to quickly handle highly similar sequences, leaving $B_{2..n}$ components to process dissimilar sequences. Sequence pairs are processed sequentially through various components until one component produces a sufficiently accurate estimation or, in the worst case, a final score-bounding component $B_n$ is reached. This $B_n$ component executes a more time-consuming algorithm but produces an exact alignment score. As a result, this cascading approach balances computational efficiency and accuracy, attending to the diverse characteristics of input sequences.

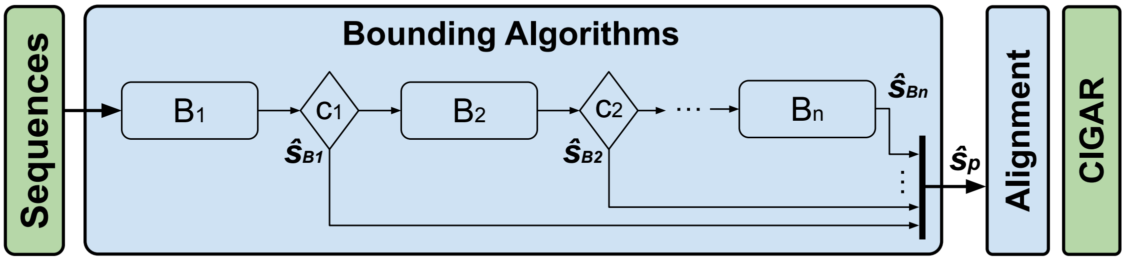


Figure S2: Block diagram of n-components cascade of score-bounding algorithms.

#### S3 WindowEd Parameter Exploration

We have performed a design space exploration of the window and overlap size parameters of the WindowEd algorithm to select the configuration of the WindowEd(S) and WindowEd(L) components used in QuickEd. We have evaluated the execution time and recall of the different configurations of $W$ window size and $O$ overlap. It is worth noting that QuickEd implements a vectorized implementation of the smallest WindowEd configuration (i.e., $W = 128$ and $O = 64$). Figure S3 shows the accuracy and execution time of each WindowEd configuration when aligning real datasets.

The analysis shows that the smaller WindowEd configuration ($W = 128$, $O = 64$) delivers the highest performance, being $1.7 - 2.1\times$ faster than the next smallest configuration ($W = 192$, $O = 64$), while only sacrificing 9% of recall in the worst-case. Consequently, we choose to utilize the $W = 128$ and $O = 64$ configuration as the WindowEd(S) component to efficiently score-bound highly-similar sequences. For the WindowEd(L) component, we choose a window size of $W = 640$ with an overlap $O = 128$. In practice, this configuration shows a good balance between execution time and accuracy, minimizing the execution time of the WindowEd(L) while reducing the number of sequences processed by the exponential BandEd (i.e., last component).

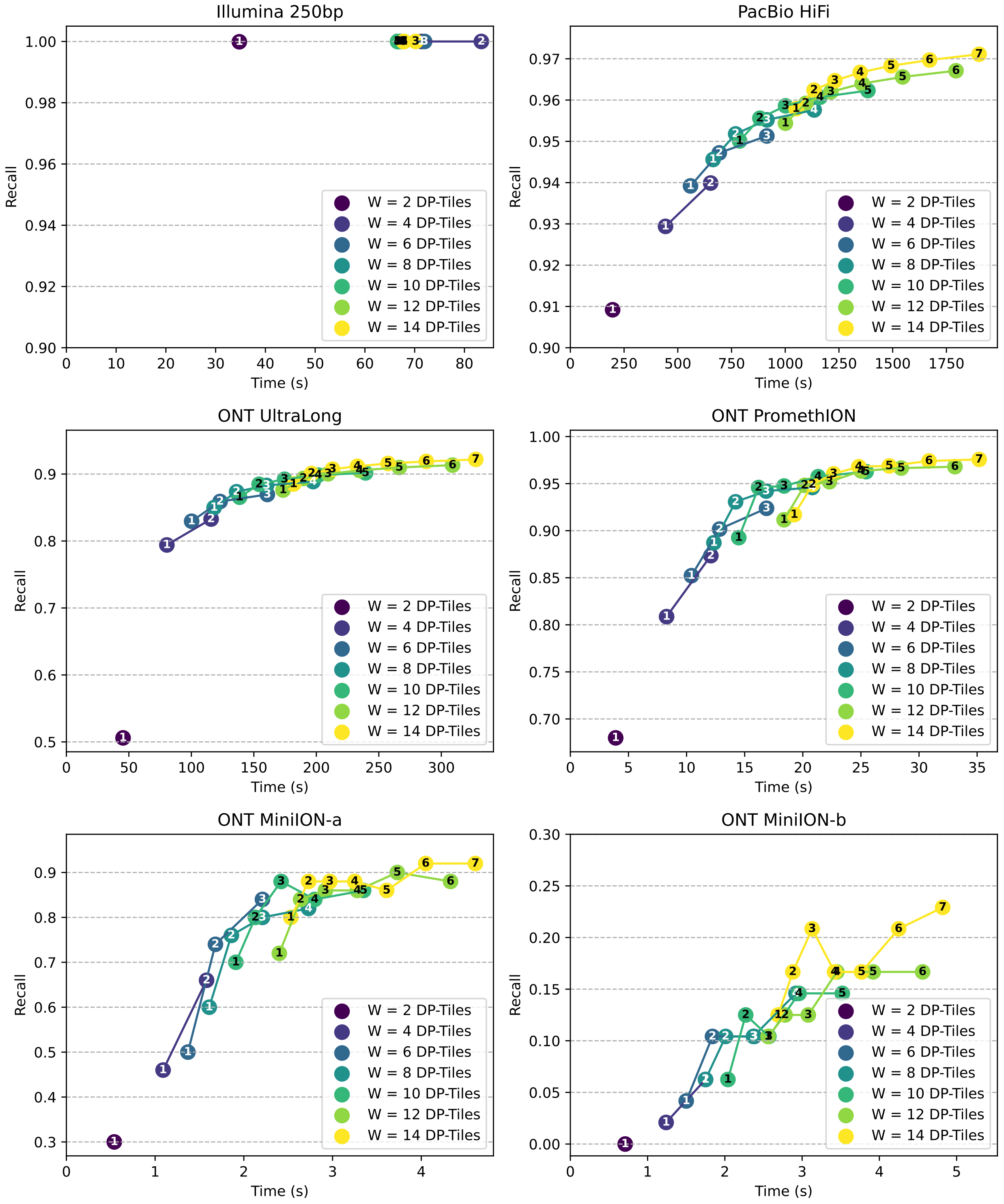


Figure S3: Design space exploration of the $W$ window size and $O$ overlap size parameters of the WindowEd algorithm. The $W$ and $O$ sizes are expressed in blocks of 64x64 DP-cells. The number inside the markers (ⓧ) represents the overlap size in blocks.

#### S4 Experimental Evaluation of Exact Algorithms using Simulated Datasets

Table S2: Time (in seconds) and memory usage (in megabytes) for the different exact alignment tools aligning simulated and real datasets. Executions taking more than 48 hours are labelled as "to" (time out) and executions as run out of memory as "oom".

| | | Time (s) | | | | | | Memory (MB) | | | | | |
|---|---|---|---|---|---|---|---|---|---|---|---|---|---|
| | | KSW2 | WFA | BiWFA | A* | Edlib | QuickEd | KSW2 | WFA | BiWFA | A* | Edlib | QuickEd |
| 10 Kbp | 1% | 1876.2 | 0.5 | 0.7 | 43.7 | 50.5 | 5.2 | 191.7 | 3.2 | 1.7 | 3.9 | 17.9 | 22.0 |
| | 5% | 18837.6 | 3.4 | 4.3 | 47.2 | 95.4 | 15.5 | 19088.0 | 28.0 | 5.2 | 22.1 | 19.1 | 30.1 |
| | 10% | oom | 18.1 | 22.3 | 69.6 | 238.2 | 69.0 | oom | 129.3 | 21.6 | 97.7 | 21.7 | 36.4 |
| | 20% | oom | 45.5 | 48.7 | 72.0 | 396.0 | 133.8 | oom | 429.7 | 28.6 | 194.9 | 26.6 | 44.8 |
| 100 Kbp | 1% | 1868.4 | 7.1 | 8.9 | 39.4 | 59.6 | 7.1 | 191.7 | 5.3 | 6.5 | 3.9 | 19.3 | 22.4 |
| | 5% | 18839.4 | 71.4 | 94.8 | 44.1 | 227.4 | 60.6 | 19101.8 | 126.9 | 7.3 | 22.2 | 20.5 | 32.9 |
| | 10% | oom | 367.8 | 466.2 | 63.6 | 714.0 | 304.8 | oom | 2186.8 | 27.2 | 92.3 | 24.2 | 39.6 |
| | 20% | oom | 755.4 | 939.0 | 67.2 | 1345.8 | 610.2 | oom | 8920.1 | 28.7 | 183.8 | 29.5 | 46.4 |
| 500 Kbp | 1% | 1873.8 | 24.0 | 32.8 | 54.8 | 70.2 | 9.4 | 192.2 | 25.3 | 3.9 | 3.8 | 19.5 | 23.4 |
| | 5% | 18839.4 | 246.0 | 327.6 | 63.6 | 375.0 | 108.0 | 19104.1 | 351.8 | 9.6 | 20.8 | 20.4 | 39.0 |
| | 10% | oom | 1257.0 | 1641.0 | 87.6 | 1332.0 | 588.6 | oom | 7871.4 | 21.8 | 84.6 | 25.3 | 34.0 |
| | 20% | oom | 2535.6 | 3333.6 | 94.2 | 2581.2 | 1173.0 | oom | 31935.6 | 43.8 | 168.4 | 30.0 | 46.5 |
| 1 Mbp | 1% | 1873.8 | 74.4 | 109.2 | 3081.0 | 89.4 | 13.2 | 193.1 | 69.5 | 3.9 | 70.2 | 19.5 | 27.6 |
| | 5% | 18873.0 | 754.8 | 1078.2 | 44994.6 | 654.6 | 212.4 | 19115.9 | 1013.4 | 9.1 | 3306.7 | 20.6 | 30.9 |
| | 10% | oom | 3807.6 | 5365.8 | oom | 2505.6 | 1080.6 | oom | 25438.9 | 26.2 | oom | 25.1 | 36.5 |
| | 20% | oom | oom | 10807.8 | oom | 4926.6 | 2165.4 | oom | oom | 42.0 | oom | 30.5 | 48.2 |
| Illumina 250bp | | 1185.0 | 7.6 | 15.8 | 1124.4 | 403.2 | 124.2 | 1.1 | 1.1 | 1.1 | 1.8 | 18.8 | 18.6 |
| PacBio HiFi | | to | 113.4 | 153.6 | 18140.4 | 5893.8 | 800.4 | to | 85.9 | 3.0 | 246.8 | 20.5 | 40.6 |
| ONT UL | | oom | oom | 7075.8 | oom | 5998.8 | 2795.4 | oom | oom | 37.0 | oom | 29.0 | 48.2 |
| ONT PromethION | | oom | 696.6 | 880.8 | 27288.6 | 884.4 | 346.2 | oom | 10299.9 | 12.6 | 52279.0 | 21.8 | 40.9 |
| ONT UUL | | oom | 316.2 | 366.0 | oom | 322.8 | 159.0 | oom | 47169.3 | 24.3 | oom | 25.5 | 43.4 |
| ONT MiniION | | oom | oom | 547.2 | oom | 431.4 | 265.2 | oom | oom | 20.8 | oom | 26.5 | 47.6 |

#### S5 QuickEd Execution Time Distribution along the Different Components

QuickEd's current implementation uses an alignment-score bounding algorithm composed of 3 components connected in cascade: WindowEd(S), WindowEd(L), and BandEd (i.e., an exponential band implementation). QuickEd's alignment-score bounding algorithm stops when a component's estimation is accurate enough. Hence, not all the components in the cascade are executed for every input sequence pair.

Table S3 shows the number of sequences processed by each component within QuickEd's alignment-score bounding algorithm. For the Illumina, PacBio, and ONT PromethION datasets, the WindowEd(S) component can process more than 95% of the sequences. By incorporating the WindowEd(L) component, we generate accurate estimations for 99-100% of the sequences. Consequently, utilizing the exponential BandEd component is only necessary to estimate approximately 0.6% of the PacBio sequences, notably reducing the performance overhead of this accurate but expensive component. However, the ONT UltraLong and MiniION datasets, characterized by having noisier sequences with longer gaps, require more accurate and time-consuming components (i.e., WindowEd(L) and BandEd)

Table S3: Number of sequences processed by each component. Note that all the sequences have to be processed by the WindowEd(S) component (first component in the bound step) and the QuickEd-Align algorithm (align step).

| | QuickEd-Bound | | | QuickEd-Align |
|---|---|---|---|---|
| | WindowEd(S) | WindowEd(L) | BandEd | |
| Illumina 250bp | 10M (100%) | 22.1k (0.22%) | 0 (0%) | 10M (100%) |
| PacBio HiFi | 1M (100%) | 38.0k (3.8%) | 5.9k (0.6%) | 1M (100%) |
| ONT UltraLong | 100k (100%) | 41k (41%) | 2k (2%) | 100k (100%) |
| ONT PromethION | 1312 (100%) | 59 (4.5%) | 0 (0%) | 1312 (100%) |
| ONT MiniION-a | 50 (100%) | 13 (26%) | 2 (4%) | 50 (100%) |
| ONT MiniION-b | 48 (100%) | 45 (93.8%) | 14 (29.1%) | 48 (100%) |

Figure S4 shows the time distribution of the different components inside QuickEd for the different datasets. As expected, QuickEd-Align execution time dominates. However, in the datasets with shorter sequences (Illumina and PacBio), the execution time of the WindowEd(S) bounding component is still significant. Aligning datasets containing short sequences, the number of DP-cells computed by the WindowEd(S) component is similar to the ones computed by QuickEd-Align since the band tends to be very small. In contrast, the execution time of WindowEd(S) using ONT datasets is negligible since the bandwidth used within QuickEd-Align is much larger.

The execution time of the most accurate components (i.e., BandEd and WindowEd(L)) impacts the overall execution time aligning PacBio, ONT UltraLong, and ONT MiniION datasets. This is due to the necessity of using these components to estimate alignment-scores for many sequences in these datasets. Conversely, in the Illumina and ONT PromethION datasets, where the number of sequences processed by each component is either zero or nearly zero, the impact on execution time is minimal or negligible. These results are expected after analyzing the number of sequences processed by the different components Table S3.

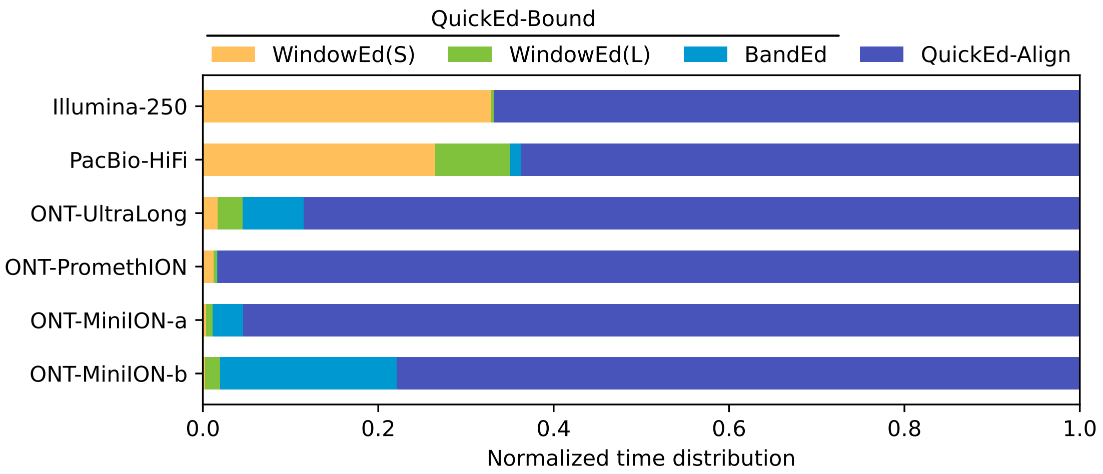


Figure S4: Normalized time distribution of the different components inside QuickEd for the different datasets.
